# Genome assembly of the hybrid grapevine *Vitis* ‘Chambourcin’

**DOI:** 10.1101/2023.01.18.524616

**Authors:** Sagar Patel, Zachary N. Harris, Jason P. Londo, Allison Miller, Anne Fennell

## Abstract

**Background:** ‘Chambourcin’ is a French-American interspecific hybrid grape variety grown in the eastern and midwestern United States and used for making wine. Currently, there are few genomic resources available for hybrid grapevines like ‘Chambourcin’.

**Results:** We assembled the genome of ‘Chambourcin’ using PacBio HiFi long-read sequencing, Bionano optical map sequencing and Illumina short read sequencing. We produced an assembly for ‘Chambourcin’ with 26 scaffolds with an N50 length of 23.3 Mb and an estimated BUSCO completeness of 97.9%. 33,791 gene models were predicted, of which 81% (27,075) were functionally annotated using Gene Ontology and KEGG pathway analysis. We identified 16,056 common orthologs between ‘Chambourcin’ gene models, *V. vinifera* ‘PN40024’ 12X.v2, VCOST.v3, Shine Muscat (*Vitis labruscana x V. vinifera*) and *V. riparia* Gloire. A total of 1,606 plant transcription factors representing 58 different gene families were identified in ‘Chambourcin’. Finally, we identified 304,571 simple sequence repeats (SSRs), repeating units of 1-6 base pairs in length in the ‘Chambourcin’ genome assembly.

**Conclusions:** We present the genome assembly, genome annotation, protein sequences and coding sequences reported for ‘Chambourcin’. The ‘Chambourcin’ genome assembly provides a valuable resource for genome comparisons, functional genomic analysis and genome-assisted breeding research.

## INTRODUCTION

Grapevines (*Vitis* spp.) represent the most economically important berry-producing plants in the world, the fruits of which are used to make wine and other beverages and are consumed as fresh or dried fruit. The European grapevine *Vitis vinifera* L. ssp. *vinifera* is believed to have been domesticated approximately 8,000 years ago from wild populations of *V. vinifera* ssp. *sylvestris* growing in western Asia and eastern Europe (Myles et al. 2011, Dong et al. 2023). Grapevine growing (viticulture) spread rapidly through Europe and the Middle East, and eventually was introduced into North America as early as the mid 1700’s and likely earlier (Pinney 1989). In addition to the introduced *V. vinifera,* North America is home to at least 20 different native *Vitis* species. Although European settlers in North America cultivated native North American *Vitis* spp.; today, few native North American grapevine species are used to make wine (e.g., *Vitis labrusca).* Despite this, many native North American *Vitis* species have become critical resources for viticulture through the development of disease resistant rootstocks (the below-ground portion of grafted vines) and hybrid scions (the above-ground portion of grafted vines) derived from interspecific hybridization between wild North American *Vitis* species and cultivated European *V. vinifera.* Hybrid derivatives of crosses between North American and European grapevine species make up a significant portion of the grapevines grown in eastern and midwestern North America, and hybrid rootstocks are used throughout most grape growing regions in the world.

*Vitis* ‘Chambourcin’ (‘Chambourcin’ from here forward) is a cultivated hybrid wine grape variety derived from crosses between North American and European *Vitis* species. ‘Chambourcin’ was developed by the private breeder Joannes Seyve in France and was introduced into the USDA-ARS repository in Geneva, NY in 1985 (Foundation Plant Services). A complex hybrid, ‘Chambourcin’ is the product of Joannes Seyve 11369 and ‘Plantet N’ that includes several North American species in its background: *V. berlandieri* Planch., *V. labrusca L., V. lincecumii* Buckley, *V. riparia* Michx., *V. rupestris* Scheele and *V. vinifera.* The full pedigree of ‘Chambourcin’ is available at https://www.vivc.de/. ‘Chambourcin’ produces black-skinned berries. Flavors of wine derived from ‘Chambourcin’ are described as black cherry, red fruit with herbaceous notes, black pepper and chocolate (winetraveler.com). ‘Chambourcin’ is grown in parts of France and Australia, as well as in Colorado, Missouri, Nebraska, New Jersey, New York, Pennsylvania, and Virginia, among others.

‘Chambourcin’ is increasing in importance as a cultivated hybrid winegrape in the central and eastern United States and it has been used in experimental rootstock vineyards aimed at understanding rootstock effects on shoot system phenotypes (Migicovsky et al. 2019; Maimaitiyiming 2020; Awale et al. 2021; Harris et al. 2021, 2022), and is the parent of new disease resistant cultivar ‘Regent’. The goals of this study were 1) to develop a high-quality reference genome for ‘Chambourcin’; and 2) to identify and annotate gene models for more accurate functional genomic analysis for this disease resistant cultivar. Work presented here advances understanding of hybrid grapevine genomics and will facilitate analyses of rootstock-scion interactions in ‘Chambourcin’ experimental vineyards.

## Methods

### PacBio HiFi, Bionano optical map and Illumina sequencing

‘Chambourcin’ leaf material was obtained from a 12-year-old experimental vineyard located at the University of Missouri Southwest Research Station in Mount Vernon, Missouri, USA. For PacBio HiFi sequencing, high molecular weight (HMW) DNA was isolated using the Nucleobond Kit (Macherey-Nagel, Bethlehem, PA) as per manufacturer’s protocol. Approximately 20 ug DNA was sheared to a center of mass of 10-20 Kb in a Megaruptor 3 system and a HiFi sequencing library was constructed following HiFi SMRTbell protocols for the Express Template Prep Kit 2.0 according to manufacturers’ recommendations (Pacific Biosciences, California). The library was sequenced using Sequel binding and sequencing chemistry v2.0 in CCS mode in a Sequel II system with movie collection (file format of HiFi data) time of 30hrs. The HiFi reads were generated with the circular consensus sequencing (CCS) mode of pbtools using a minimum Predicted Accuracy of 0.990.

For Bionano data, DNA was isolated from fresh young leaf tissue from a 12-year-old experimental vineyard located at the University of Missouri Southwest Research Station in Mount Vernon, Missouri, USA using the Prep™ Plant DNA Isolation and labeled using the Bionano Prep™ DNA Labeling Kit (Direct Label and Stain (DLS)), (Bionano Genomics, San Diego CA). In total, 500 ng uHMW DNA was used for the DLS reaction. DNA was incubated in the presence of DLE-1 Enzyme, DL-Green and DLE-1 Buffer for 3:20 h at 37 °C, followed by a proteinase K digestion at 50C for 30 minutes, double cleanup of unincorporated DL-Green label. The resulting DLS sample was combined with Flow Buffer, DTT and DNA stain, mixed at slow speed in a rotator mixer for an hour and incubated overnight at 4 °C. The labeled sample was then loaded onto a Bionano flow cell in a Saphyr System for separation, imaging, and creation of digital molecules according to the manufacturer’s recommendations (https://bionanogenomics.com/support-page/saphyr-system). The raw molecule set was filtered to a molecule length of 250 kb and minimum labels of nine CTTAAG labels per molecule. Bionano maps were assembled without pre-assembly using the non-haplotype parameters with no CMPR cut and without extend-split. Bionano software (Solve, Tools and Access, v1.5.1) was used for data visualization, processing and assembly of Bionano maps. The PacBio HiFi and Bionano sequencing were done at Corteva Agriscience, Johnston, Iowa -USA.

For Illumina whole genome data, DNA was extracted from ‘Chambourcin’ leaf tissue collected from the USDA Grape Germplasm Collection located in Geneva, New York. DNA was extracted using Qiagen DNeasy Plant Mini Kits (Qiagen, Valencia, CA, USA) and assessed for purity and concentration using a NanoDrop spectrophotometer and Qubit fluorometer. DNA was cleaned using a Qiagen Dneasy PowerClean Pro Cleanup Kit. DNA libraries were prepared and shotgun Illumina sequenced at Novogene (San Diego, CA, USA) with paired-end 150 nt reads with 40X coverage. The raw Illumina reads were trimmed with Trimmomatic (v0.39) (Bolger et al., 2014) using HEADCROP:4 MINLEN:70 parameters.

### Genome size estimation

The PacBio HiFi reads and 19 nt k-mers were used to estimate genome heterozygosity using jellyfish (v2.3.0) (Marçais et al., 2011). The resulting “.histo” file was visualized with GenomeScope (Vurture et al., 2017).

### Genome Assembly

PacBio HiFi assembly was generated using the HiFiasm assembler (v0.13-r308) (Cheng et al., 2021) with default parameters. To reduce the number of small, low-coverage artifactual contigs often generated by Hifiasm, the assembly was filtered to exclude less than 70,000 bp contigs. Resulting HiFi contigs were merged with the DLS Bionano maps with Bionano Solve (v3.5.1) using the hybridscaffold.pl script of Bionano Solve (v3.5.1) to get a hybrid assembly. Each scaffold of the hybrid assembly was then checked and small overlapping contigs were curated and removed to make a contiguous sequence. This curated diploid assembly was examined to identify alternative contigs using Purge Haplotigs (v1.1.1) (Roach et al., 2018) and the primary assembly and haplotig assemblies were created. We mapped trimmed Illumina whole genome sequences to both assemblies separately with bowtie2 (v2.3.4) and samtools (v1.9), and the resulting .bam files were used for polishing both assemblies using Pilon (v1.23) and the final assembly (primary assembly) and haplotig assemblies were prepared. In this study, we used only the primary assembly for all downstream analysis but the haplotigs are maintained to cover the total heterozygous genome. Scaffolds were aligned to the *V. vinifera* ‘PN40024’ 12X.v2 (Canaguier et al., 2017) reference genome using minimap2 (v2.17) (Heng Li et al., 2018) and renamed based on longest alignment with reference genome *V. vinifera* ‘PN40024’ 12X.v2 chromosomes. We mapped two thousand Chambourcin rhAmpSeq marker sequences to the ‘Chambourcin’ genome assembly using BWA aligner (v0.7.17) (Heng Li, et al, 2009). The rhAmpSeq markers were designed to target the core *Vitis* genome and were developed from gene rich collinear regions of 10 *Vitis* genomes (Zou, C., et al., 2020). These markers aid in mapping contig on chromosomes and checking orientation.

### Genome assembly assessment and Dotplot

All assemblies generated by PacBio HiFi and Bionano data were assessed by Benchmarking Universal Single-Copy Orthologs (BUSCO) (v5.4.2) (Simão et al., 2015) with genome mode and embryophyta_odb10 dataset. The alignment of two genomes were obtained using minimap2 (v2.17) (Heng Li et al., 2018) with default parameters where the ‘Chambourcin’ primary assembly was considered as query and *V. vinifera* ‘PN40024’ 12X.v2, Shine Muscat and *V. riparia* Gloire considered as the reference genome. The dotplot was obtained using R script (https://github.com/tpoorten/dotPlotly).

### De novo gene prediction, functional annotation and orthologous genes

De novo repeats were identified with RepeatModeler2 (v2.0.2a) (Jullien Li et al., 2020) and then repeats were masked by RepeatMasker (v4.1.1) (Smit et al., 2015). ‘Chambourcin’ RNA-seq data downloaded from previous published study (Migicovsky et al., 2019) and trimmed using Trimmomatic (v0.39) (Bolger et al., 2014) with HEADCROP:15 LEADING:30 TRAILING:30 MINLEN:20 parameters. The trimmed ‘Chambourcin’ RNA-seq reads then mapped to the masked ‘Chambourcin’ primary genome assembly using HISAT2 (v2.1.0) (Kim et al., 2019) with default parameters. The resulting alignments (.bam files) and protein sequences of the *V. vinifera* ‘PN40024’ 12X.v2, VCost.v3 (Canaguier et al., 2017) were used for gene prediction using BRAKER2 (v2.1.6) (Tomáš Brůna et al., 2021) with --prg=gth --gth2traingenes --gff3 parameters. The resulting gene predictions (proteins, coding sequences and annotations) were completed separately for the ‘Chambourcin’ primary assembly and for the ‘Chambourcin’ haplotig assembly. The quality of the predicted proteins were assessed using BUSCO (v5.4.2) (Simão et al., 2015) with protein mode and embryophyta_odb10 dataset. The predicted proteins of *Vitis* ‘Chambourcin’ primary assembly were then functionally annotated using EggNOG-mapper (http://eggnog-mapper.embl.de/) and related Gene Ontology (GO), KEGG pathway and other functional information. The Gene Ontology plot was developed using WEGO tool (Ye J, 2018). For Orthologous gene models analysis, the sequences of ‘Chambourcin’ primary gene models, *V. vinifera* PN40024 12X.v2, VCost.v3 (Canaguier et al., 2017), Shine Muscat (*Vitis labruscana x V. vinifera;* Shirasawa et al., 2022) and *V. riparia* Gloire (Girollet et al., 2020) were analyzed using OrthoVenn2 (Ling Xu et al., 2019) (https://orthovenn2.bioinfotoolkits.net/home) with default settings using E-value: 1e-5 and inflation value: 1.5.

### Plant transcription factors prediction, phylogenetic tree and WRKY classification

The plant transcription factors for ‘Chambourcin*’* primary assembly gene models and *V. vinifera* PN40024 12X.v2, VCost.v3 gene models were identified using PlantTFDB (5.0) (Jin JP et al., 2017) (http://planttfdb.gao-lab.org/). The identified transcription factors were divided into subfamilies according to their sequence relationship with *V. vinifera*. For circular phylogenetic tree and WRKY classification, WRKY sequences of ‘Chambourcin*’* primary gene models and *V. vinifera* PN40024 12X.v2, VCost.v3 gene models retrieved from PlantTFDB (5.0) (Jin JP et al., 2017) (http://planttfdb.gao-lab.org/) and aligned using ClustalW method in MEGA7 (Kumar et al., 2016). Phylogenetic analysis was carried out using the Neighbor-Joining method with 1000 bootstrap replications and the evolutionary distances were computed using the Poisson correction method with Pairwise Deletion option. WRKY classification of ‘Chambourcin*’* primary gene models carried out using the same method described in (Patel et al., 2020).

### Synteny and Simple Sequence Repeats (SSRs)

The ‘Chambourcin’ masked primary genome assembly and gene annotations were aligned to *V. vinifera* ‘PN40024’ 12X.v2 (Canaguier et al., 2017), Shine Muscat (Shirasawa et al., 2022) and *V. riparia* ‘Gloire’ (Girollet et al., 2020) genomes and gene annotations separately using the ‘promer’ option of the MUMmer program in SyMAP (v4.2) (C. Soderlund et al., 2010). We employed MIcroSAtellite (MISA) (Sebastian Beier et al., 2017) (https://webblast.ipk-gatersleben.de/misa/) to find Simple Sequence Repeats (SSRs) in unmasked ‘Chambourcin’ primary genome assembly.

## Results and Discussion

### Genome Sequencing and Assembly of ‘Chambourcin’

We generated a high quality and contiguous genome sequence of ‘Chambourcin’ using PacBio HiFi Sequencing, Bionano third-generation DNA sequencing and Illumina short read sequencing. A total of 1,634,814 PacBio HiFi filtered reads were produced with an average length of 16,148 bp and genome coverage of 28X. The filtered Bionano data resulted in a subset of 1,243,428 molecules with a total length of 429,808.857 (Mbp) and coverage of 188.70X. In total, 124 Bionano maps with a total length of 962.964 (Mbp) and an N50 of 13,725 bp were assembled, corresponding to the diploid complement. A total of 154,152,068 filtered Illumina short reads and genome coverage of 40X were generated for genome polishing. We estimated heterozygosity to be 2.28% in the ‘Chambourcin’ genome (Figure 1), which is higher than estimates for heterozygosity in any of the other *Vitis* genomes sequenced to date (Canaguier et al., 2017; Girollet et al., 2020; Patel et al., 2020). Relatively higher levels of heterozygosity in the ‘Chambourcin’ compared to other *Vitis* species are expected given the complex interspecific pedigree of this cultivar. A GenomeScope plot of clean reads demonstrated two peaks of coverage; the first peak located at 25X coverage corresponds to the heterozygous portion of the genome, and the second peak at 52X coverage corresponds to the homozygous portion of the genome (Figure 1).

**Figure 1.**
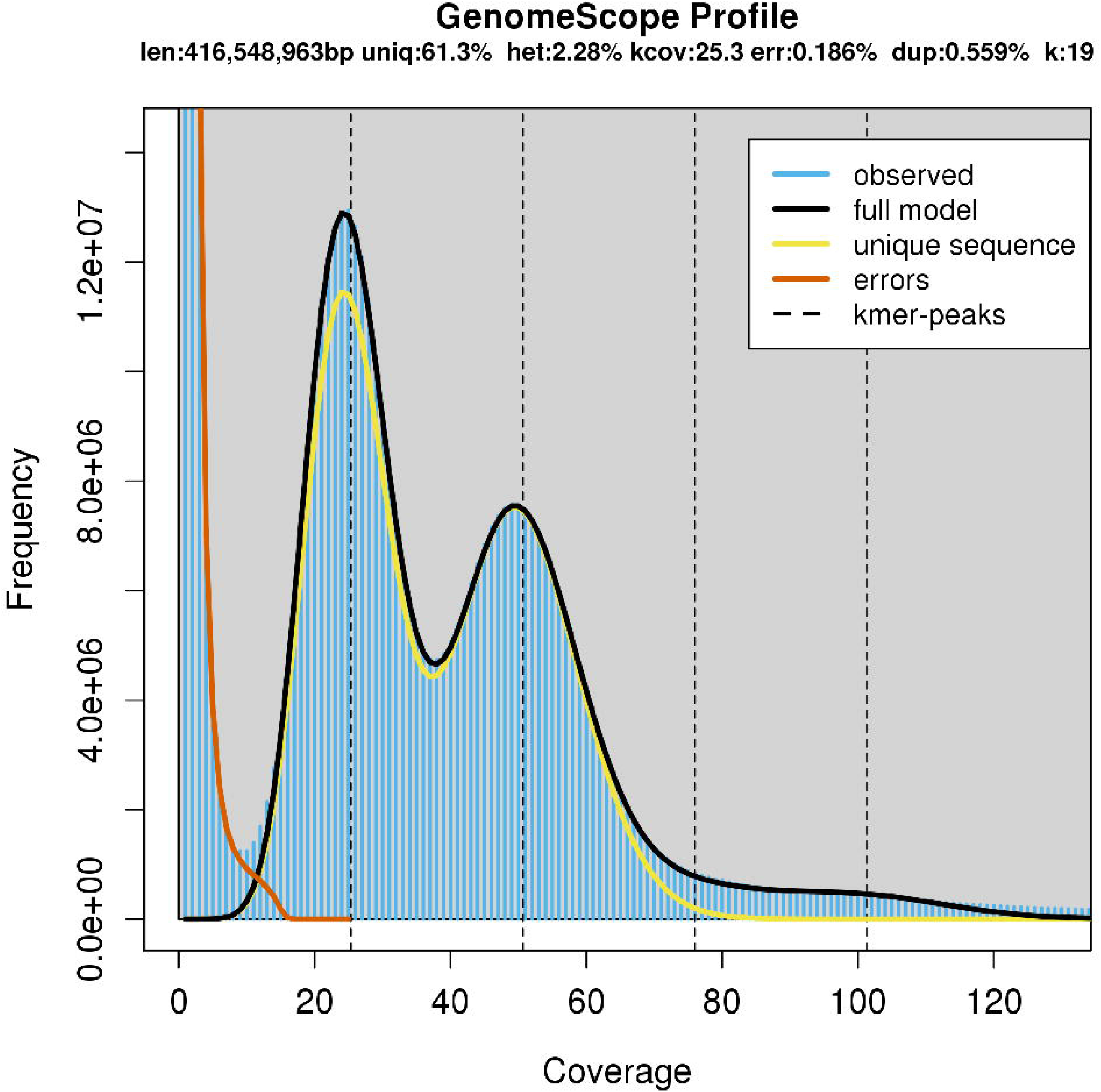
GenomeScope plot estimating heterozygosity of the *Vitis* ‘Chambourcin’.

A de novo ‘Chambourcin’ genome was assembled using HiFi data, Bionano data and Illumina data. First, a contig assembly of the PacBio HiFi reads resolved the reads into 196 contigs with an N50 of 12,215,205 bp and total length of 949,347,381 bp (Table 1). The PacBio HiFi contig assembly was then merged with the Bionano maps to get an initial hybrid assembly comprising 67 scaffolds with a N50 length of 16,400,326 bp, a maximum scaffold length of 39,458,994 bp, and total scaffold lengths of 903,810,753 bp (Table 1). After manual curation, the hybrid assembly included 64 scaffolds with an N50 length of 16,278,793 bp, maximum length of 39,458,994 bp and total length of 869,222,201 bp (Table 1). The hybrid assembly was partitioned into a final primary assembly (493,554,689 bp) and a haplotig assembly (375,458,233 bp) (Table 1). The final primary assembly after polishing for ‘Chambourcin’ contained 26 scaffolds with an N50 length of 23,325,629 bp and a longest scaffold length of 39,456,434 bp (Table 1). The secondary haplotig assembly after polishing contained 38 haplotig scaffolds with a N50 length of 1,2462,019 bp and a longest scaffold length of 28,439,729 bp (Table 1). We identified 97.9% Complete BUSCOs (C) for primary genome assembly and 73.1% Complete BUSCOs (C) for haplotig genome assembly (Table 1).

**Table 1.**
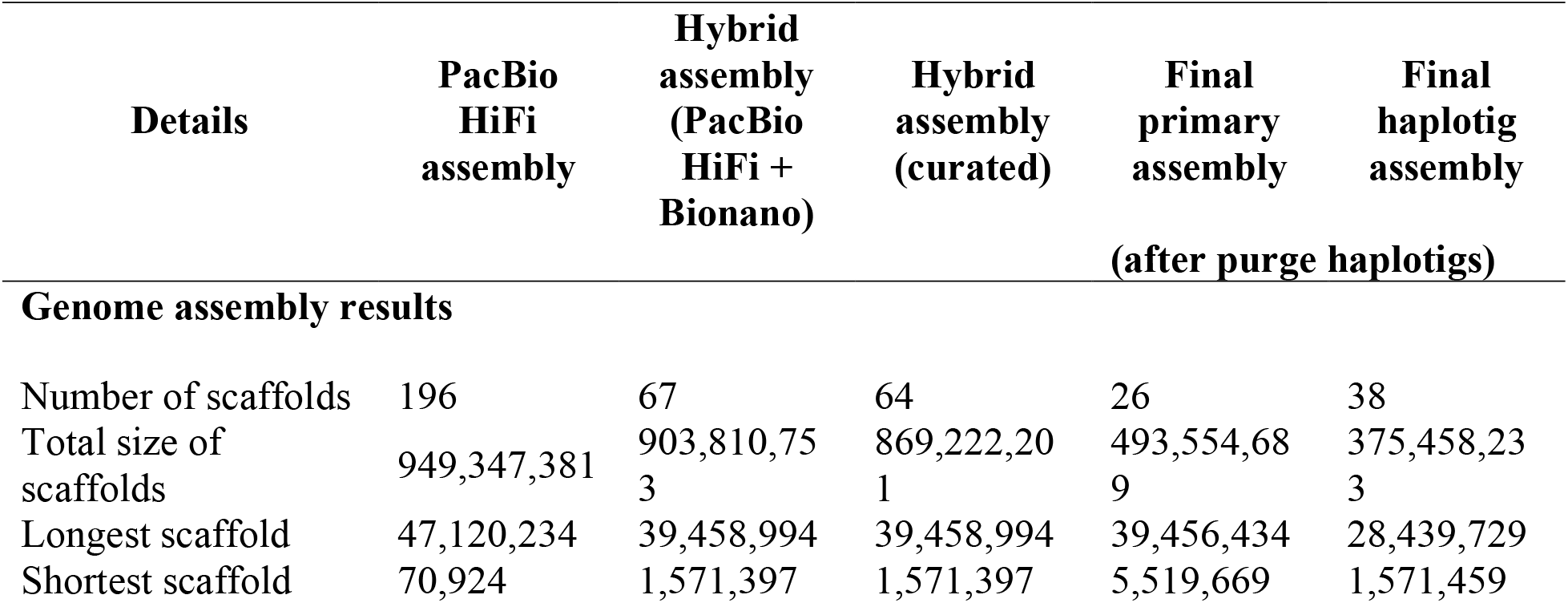
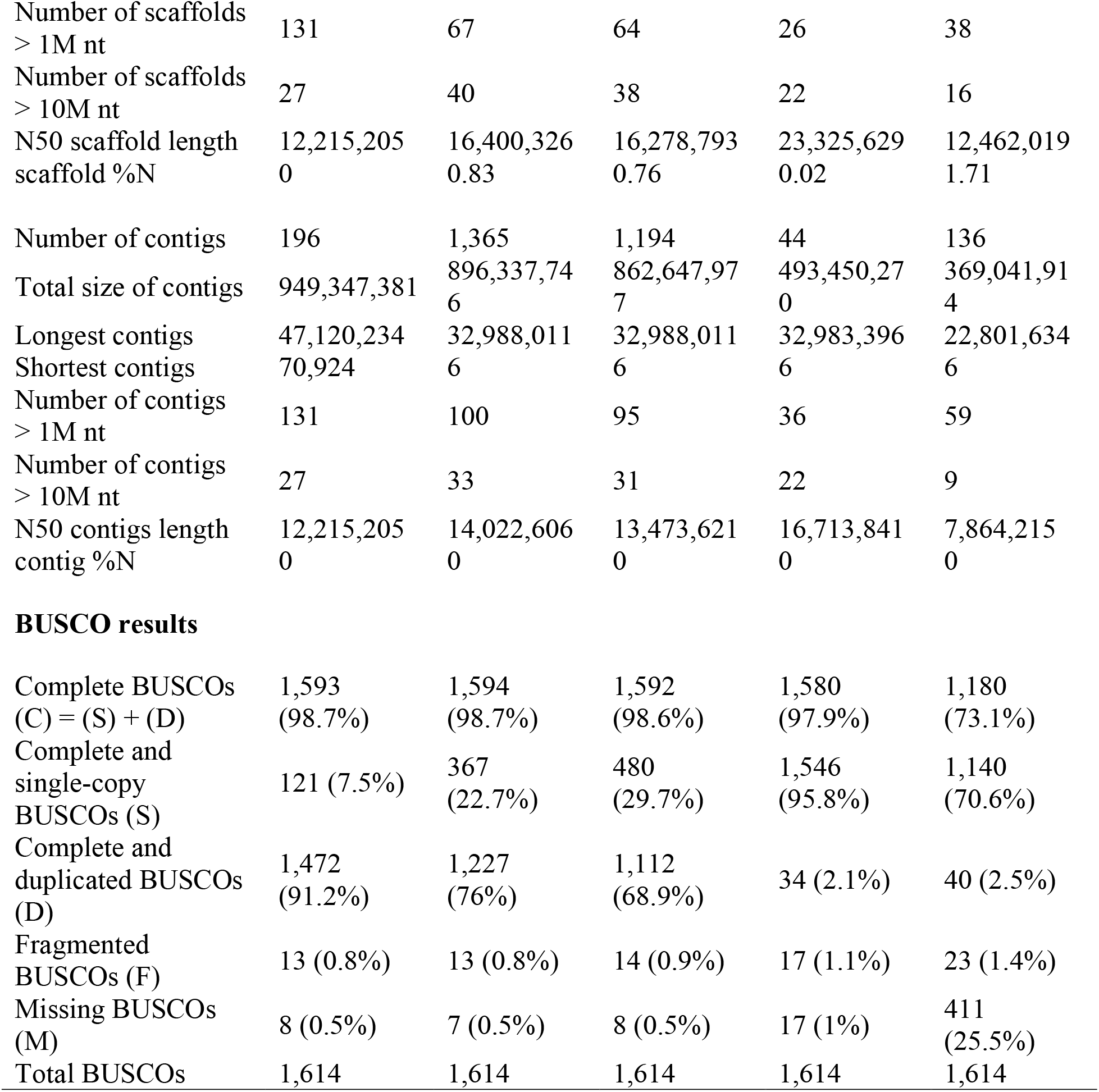
‘Chambourcin’ genome assembly descriptive statistics and BUSCO results.

| Details | PacBio<br>HiFi<br>assembly | Hybrid<br>assembly<br>(PacBio<br>HiFi +<br>Bionano) | Hybrid<br>assembly<br>(curated) | Final<br>primary<br>assembly<br>(after purge haplotigs) | Final<br>haplotig<br>assembly |
| --- | --- | --- | --- | --- | --- |
| **Genome assembly results** |  |  |  |  |  |
| Number of scaffolds | 196 | 67 | 64 | 26 | 38 |
| Total size of<br>scaffolds | 949,347,381 | 903,810,753 | 869,222,201 | 493,554,689 | 375,458,233 |
| Longest scaffold | 47,120,234 | 39,458,994 | 39,458,994 | 39,456,434 | 28,439,729 |
| Shortest scaffold | 70,924 | 1,571,397 | 1,571,397 | 5,519,669 | 1,571,459 |
| Number of scaffolds > 1M nt | 131 | 67 | 64 | 26 | 38 |
| Number of scaffolds > 10M nt | 27 | 40 | 38 | 22 | 16 |
| N50 scaffold length | 12,215,205 | 16,400,326 | 16,278,793 | 23,325,629 | 12,462,019 |
| scaffold %N | 0 | 0.83 | 0.76 | 0.02 | 1.71 |
| Number of contigs | 196 | 1,365 | 1,194 | 44 | 136 |
| Total size of contigs | 949,347,381 | 896,337,746 | 862,647,977 | 493,450,270 | 369,041,914 |
| Longest contigs | 47,120,234 | 32,988,011 | 32,988,011 | 32,983,396 | 22,801,634 |
| Shortest contigs | 70,924 | 6 | 6 | 6 | 6 |
| Number of contigs > 1M nt | 131 | 100 | 95 | 36 | 59 |
| Number of contigs > 10M nt | 27 | 33 | 31 | 22 | 9 |
| N50 contigs length | 12,215,205 | 14,022,606 | 13,473,621 | 16,713,841 | 7,864,215 |
| contig %N | 0 | 0 | 0 | 0 | 0 |
| BUSCO results |  |  |  |  |  |
| Complete BUSCOs (C) = (S) + (D) | 1,593 (98.7%) | 1,594 (98.7%) | 1,592 (98.6%) | 1,580 (97.9%) | 1,180 (73.1%) |
| Complete and single-copy BUSCOs (S) | 121 (7.5%) | 367 (22.7%) | 480 (29.7%) | 1,546 (95.8%) | 1,140 (70.6%) |
| Complete and duplicated BUSCOs (D) | 1,472 (91.2%) | 1,227 (76%) | 1,112 (68.9%) | 34 (2.1%) | 40 (2.5%) |
| Fragmented BUSCOs (F) | 13 (0.8%) | 13 (0.8%) | 14 (0.9%) | 17 (1.1%) | 23 (1.4%) |
| Missing BUSCOs (M) | 8 (0.5%) | 7 (0.5%) | 8 (0.5%) | 17 (1%) | 411 (25.5%) |
| Total BUSCOs | 1,614 | 1,614 | 1,614 | 1,614 | 1,614 |

The ‘Chambourcin’ primary genome assembly was aligned to the reference genome *V. vinifera* ‘PN40024’ 12X.v2 (Canaguier et al., 2017) (cite GigaDB Table 1 here), Shine Muscat (Shirasawa et al., 2022) and *V. riparia* ‘Gloire’ (Girollet et al., 2020) and a dot-plot was generated to facilitate comparisons among genomes. Collinearity between ‘Chambourcin’ and *V. vinifera* ‘PN40024’ 12X.v2, Shine Muscat and *V. riparia* ‘Gloire’ was observed as a straight diagonal line without large gaps in the dot plot, confirming high synteny of the ‘Chambourcin’ genome with *V. vinifera* ‘PN40024’ 12X.v2 (Figure 1A), Shine Muscat (Figure 1B) and *V. riparia* ‘Gloire’ (Figure 1C). To further validate the Chambourcin’ genome assembly, we mapped Chambourcin rhAmpSeq markers to the ‘Chambourcin’ genome assembly and found 99% of rhAmpSeq markers mapped to ‘Chambourcin’ scaffolds and mapped to the same chromosomes and positions that the markers were derived from in the collinear *Vitis* core genome (cite GigaDB Table 2 here).

Synteny analyses of the ‘Chambourcin’ primary genome assembly with *V. vinifera* ‘PN40024’ 12X.v2, Shine Muscat and *V. riparia* ‘Gloire’ genomes were used to identify syntenic blocks between species. The ‘Chambourcin’ primary assembly scaffolds aligned with larger syntenic blocks and covered the whole chromosomes of *V. vinifera* PN40024 12X.v2 (Figure 1D and Figure 1E), Shine Muscat (Figure 1D) and *V. riparia* ‘Gloire’ (Figure 1E). This alignment of the primary genome with *V. vinifera* PN40024 12X.v2, Shine Muscat and *V. riparia* ‘Gloire’ indicated highly contiguous ‘Chambourcin’ scaffolds useful for a comparative genomic analysis.

### Repeat Sequence Annotation

Repeated regions were binned into seven different classes: long interspersed nuclear elements (LINEs) (4.43%), long terminal repeats (LTRs) (15.66%), DNA transposons (2.03%), rolling-circles (0.58%), low complexity repeats (0.37%), simple repeats (1.21%) and unclassified repeats (31.95%) (Table 2). The repetitive sequence content in the ‘Chambourcin’ primary genome assembly (56.23%) was higher than previously reported for *V. riparia* ‘Manitoba 37’ (46%) (Patel et al., 2020), *V. vinifera* ‘PN40024’ 12X.v2 (35.12%) (Canaguier et al., 2017), Shine Muscat (48%) (Shirasawa et al., 2022) and *V. riparia* (33.94%) ‘Gloire’ (Girollet et al., 2020). Simple sequence repeats (SSRs) are tandem repeats of DNA that have been used to develop robust genetic markers. We identified 304,571 simple sequence repeats (SSRs), repeating units of 1-6 base pairs in length, in the ‘Chambourcin’ primary genome assembly (Figure 2; cite GigaDB Table 3 here).

**Table 2.** Repetitive sequences in the ‘Chambourcin*’* genome assembly.

| Details | Primary assembly | Haplotig assembly |
| --- | --- | --- |
| Total length: | 493,554,689 bp | 375,458,233 bp |
| Bases masked: | 277,503,542 bp (56.23%) | 211,395,579 bp (56.30%) |
| Retroelements: | 99,122,846 bp (20.08%) | 82,725,835 bp (22.03%) |
| LINES: | 21,852,466 bp (4.43%) | 15,553,322 bp (4.14%) |
| RTE/Bov-B | 180,809 bp (0.04%) | 133,053 bp (0.04%) |
| L1/CIN4 | 21,671,657 bp (4.39%) | 15,420,269 bp (4.11%) |
| LTR elements: | 77,270,380 bp (15.66%) | 67,172,513 bp (17.89%) |
| Ty1/Copia | 39,408,750 bp (7.98%) | 32,756,463 bp (8.72%) |
| Gypsy/DIRS1 | 33,756,474 bp (6.84%) | 31,170,356 bp (8.30%) |
| DNA transposons | 9,995,420 bp (2.03%) | 7,711,872 bp (2.05%) |
| hobo-Activator | 3,826,157 bp (0.78%) | 2,619,712 bp (0.70%) |
| Tourist/Harbinger | 640,576 bp (0.13%) | 454,161 bp (0.12%) |
| Rolling-circles | 2,864,922 bp (0.58%) | 2,386,306 bp (0.64%) |
| Unclassified: | 157,704,064 bp (31.95%) | 112,960,798 bp (30.09%) |
| Total interspersed repeats: | 266,822,330 bp (54.06%) | 203,398,505 bp (54.17%) |
| Simple repeats: | 5,980,823 bp (1.21%) | 4,443,253 bp (1.18%) |
| Low complexity: | 1,835,467 bp (0.37%) | 1,167,515 bp (0.31%) |

**Figure 2.**
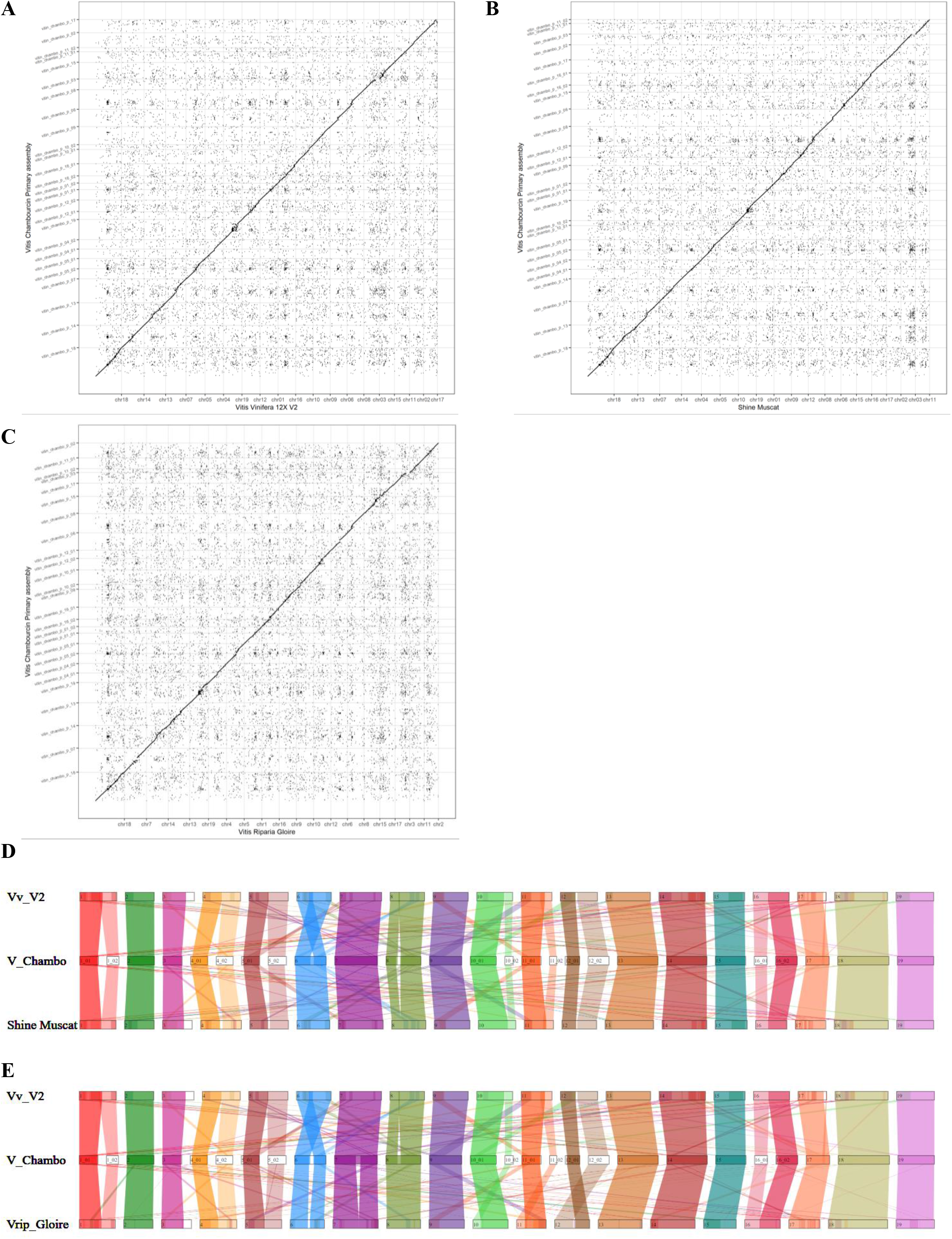
Comparative study of the ‘Chambourcin’ genome assembly. (A) Dotplot of the ‘Chambourcin’ primary genome assembly and *V. vinifera* ‘PN40024’ 12X.v2. (B) Dotplot of the ‘Chambourcin’ primary genome assembly and Shine Muscat. (C) Dotplot of the ‘Chambourcin’ primary genome assembly and *V. riparia* Gloire (D) Synteny between ‘Chambourcin’ primary genome assembly, *V. vinifera* PN40024 12X.v2 genome and Shine Muscat. (E) Synteny between ‘Chambourcin’ primary genome assembly, *V. vinifera* PN40024 12X.v2 genome and *V. riparia* Gloire genome.

### Gene Annotation and orthologous genes

A total of 33,791 gene models were predicted for the ‘Chambourcin’ primary genome assembly. We identified 94.6% complete BUSCOs (C); of these 86.9% were designated single-copy BUSCOs (S) and 7.7% were designated duplicated BUSCOs (D) (Table 3). As evidenced by the high number of complete single-copy genes identified, the BUSCO results indicate that the ‘Chambourcin’ primary genome assembly offers comprehensive coverage of expected gene space. Functional annotation of the ‘Chambourcin’ primary gene models (33,791) was carried out using the EggNOG database (http://eggnog-mapper.embl.de/) (cite GigaDB Table 4 here). In total 27,075 ‘Chambourcin’ primary proteins were annotated and 86% (22,977) of the ‘Chambourcin’ primary proteins annotated with *V*. *vinifera* V1 gene models (cite GigaDB Table 4 here). Out of the total 27,075 ‘Chambourcin’ annotated primary proteins, 13,311 gene models were identified with Gene Ontology (GO) accessions (cite GigaDB Table 4 here) and further classified into three sub-ontologies: biological process (11,399), cellular component (11,472) and molecular function (9,977) (Figure 4F) (cite GigaDB Table 4 here). A total of 8,460 ‘Chambourcin’ primary proteins were annotated with KEGG pathway IDs (cite GigaDB Table 4 here). Using OrthoVenn2 (https://orthovenn2.bioinfotoolkits.net/home), we identified a total of 16,056 common orthologs between ‘Chambourcin’ primary gene models, *V. vinifera* PN40024 12X.v2 annotation, Shine Muscat and *V. riparia* ‘Gloire’ (Figure 4A). In total, 16,476 orthologous gene models were found between the ‘Chambourcin’, Shine Muscat and *V. riparia ‘*Gloire’ (Figure 4B), while 19,477 gene models were orthologous with *V. vinifera* PN40024 12X.v2 VCost.v3 proteins (Figure 4C) and 18,669 gene models were orthologous with Shine Muscat (Figure 4E) and 18,183 gene models were orthologous with *V. riparia* ‘Gloire’ (Figure 4D).

**Table 3.** ‘Chambourcin’ gene prediction and BUSCO results of protein sequences.

| Details | Primary assembly | Haplotig assembly |
| --- | --- | --- |
| **Gene prediction results** |  |  |
| Total CDS and protein | 33,791 | 24,018 |
| Total CDS (bp) | 36,761,139 | 25,814,082 |
| Mean CDS (bp) | 1,087.9 | 1,074.8 |
| Longest CDS (bp) | 15,867 | 21,819 |
| Total protein (bp) | 12,219,929 | 8,580,681 |
| Mean protein (bp) | 361.6 | 357.3 |
| Longest protein (bp) | 5,288 | 7,272 |
| **BUSCO results** |  |  |
| Complete BUSCOs (C) = (S) + (D) | 1,528 (94.6%) | 1,136 (70.3%) |
| Complete and single-copy BUSCOs (S) | 1,403 (86.9%) | 1,032 (63.9%) |
| Complete and duplicated BUSCOs (D) | 125 (7.7%) | 104 (6.4%) |
| Fragmented BUSCOs (F) | 50 (3.1%) | 31 (1.9%) |
| Missing BUSCOs (M) | 36 (2.3%) | 447 (27.8%) |
| Total BUSCOs | 1,614 | 1,614 |

**Figure 3.**
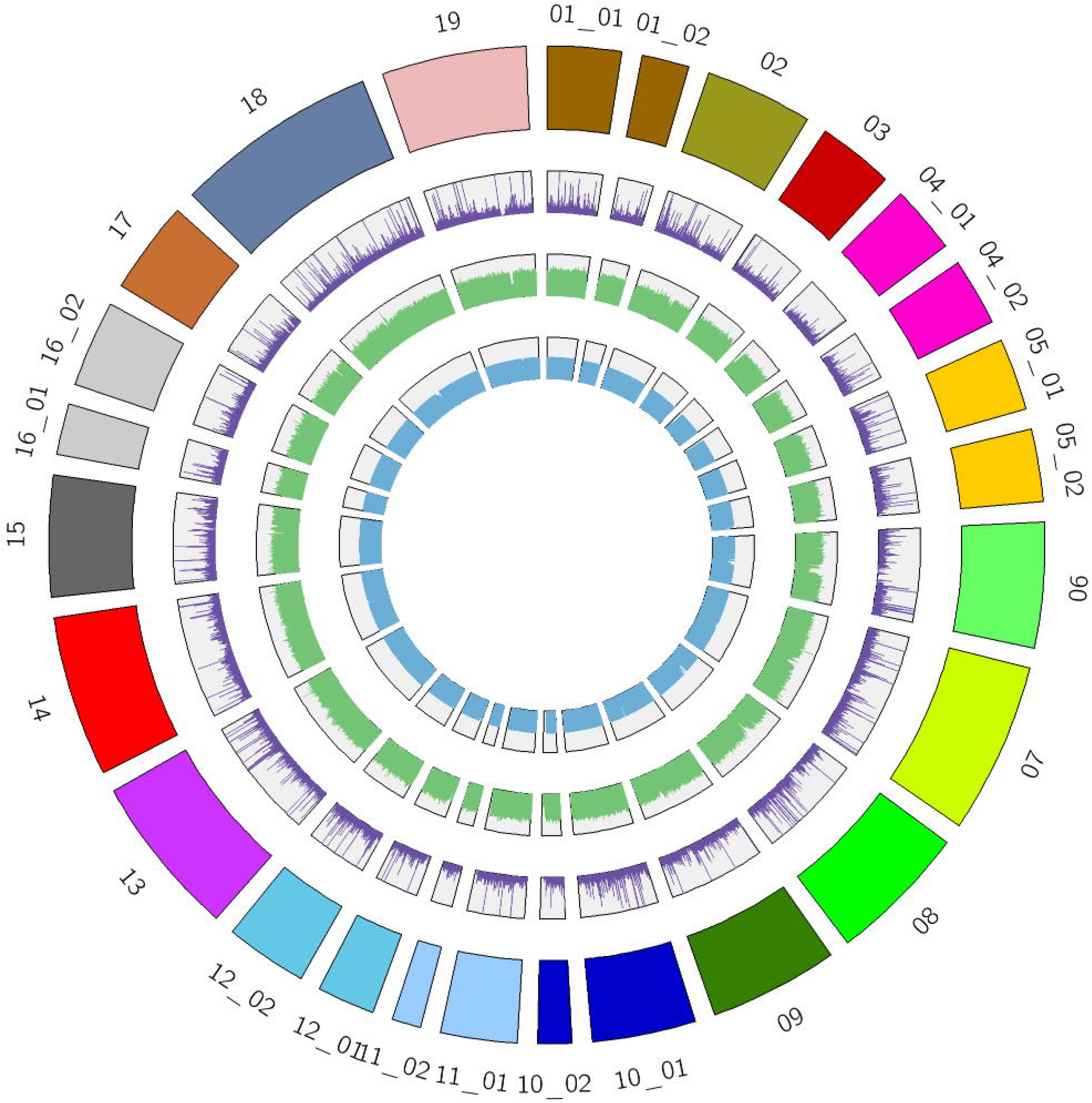
Circos plot. Outer ring represents all scaffolds of ‘Chambourcin’ primary genome assembly in different colors. The second ring of purple color represents the Simple Sequence Repeats (SSRs). The third ring of green color represents the Repetitive sequences and the fourth ring of blue color represents gene annotations.

**Figure 4.**
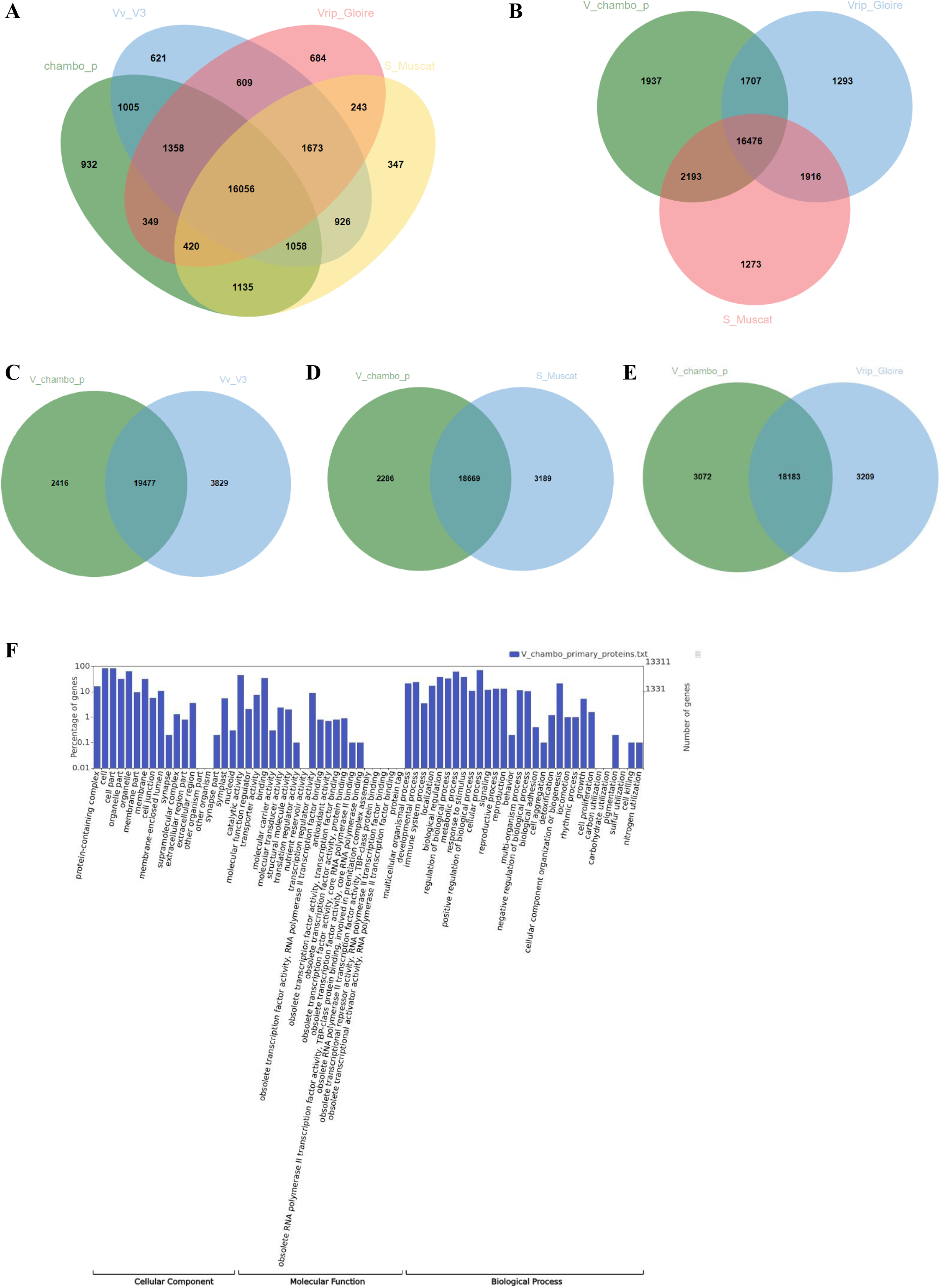
Venn diagram of ‘Chambourcin’ primary proteins with other grapevine species. (A) Venn diagram of Orthologous genes in the ‘Chambourcin’ primary proteins, *V. vinifera* PN40024 12X.v2, VCost.v3, Shine Muscat and *V. riparia* Gloire. (B) Orthologous genes in the ‘Chambourcin’ primary proteins, Shine Muscat and *V. riparia* Gloire. (C) Orthologous genes in ‘Chambourcin’ primary proteins and *V. vinifera* PN40024 12X.V3 proteins. (D) Orthologous genes in ‘Chambourcin’ primary proteins and Shine Muscat. (E) Orthologous genes in ‘Chambourcin’ primary proteins and *V. riparia* Gloire species. (F) Gene Ontology result for ‘Chambourcin’ primary proteins.

### Plant transcription factors and Chambourcin WRKY transcription factor classification

Using the Plant Transcription Factor Database (PlantTFDB 5.0 - http://planttfdb.gao-lab.org/), a total of 1,606 plant transcription factors representing 58 different gene families were identified from ‘Chambourcin’ primary proteins (cite GigaDB Table 5 here). A similar number of transcription factors were identified for the AP2, NAC, RAV and WRKY gene families as found in *V. vinifera* ‘ PN40024’ 12X.v2, VCost.v3 (cite GigaDB Table 5 here). There were 65 WRKY sequences identified in ‘Chambourcin’ and 62 in *V. vinifera* PN40024 12X.v2, VCost.v3 (Figure 5) (Table 4; cite GigaDB Table 5 here). WRKY transcription factors regulate many processes in plants and algae, such as the responses to biotic and abiotic stresses and seed dormancy. The Chambourcin WRKY subfamily classification was similar to *V. vinifera* ‘PN40024’ 12X.v2 and *V. riparia* ‘Manitoba 37’ (Table 4). These results show the high coverage of ‘Chambourcin’ primary proteins.

**Figure 5.**
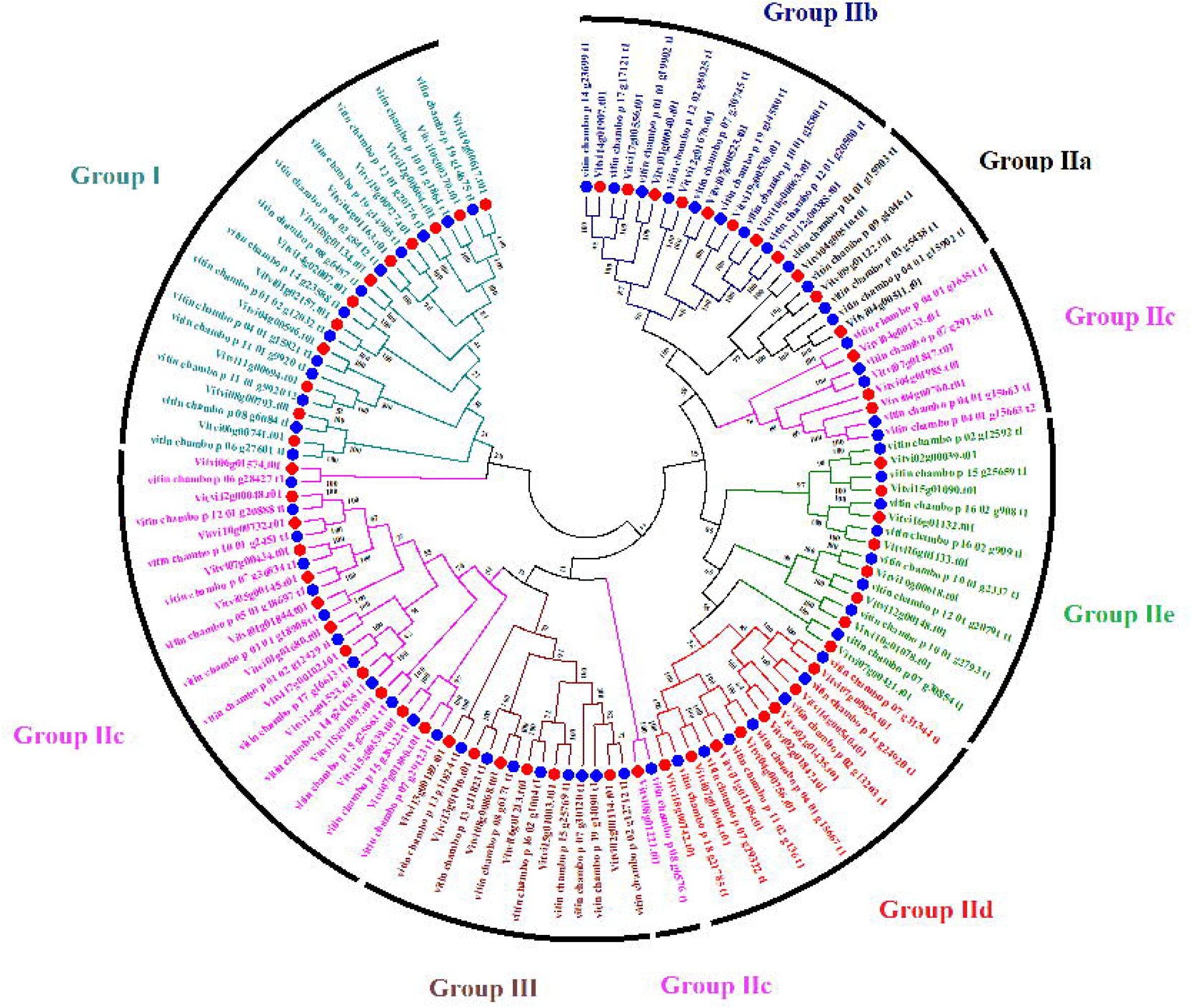
‘Chambourcin’ and *V. vinifera* PN40024 12X.v2, VCost.v3 WRKY transcription factors. Blue dots represent ‘Chambourcin’ and red dots represent *V. vinifera* PN40024 12X.v2, VCost.v3. Bootstrap values displayed are at nodes.

**Table 4.**
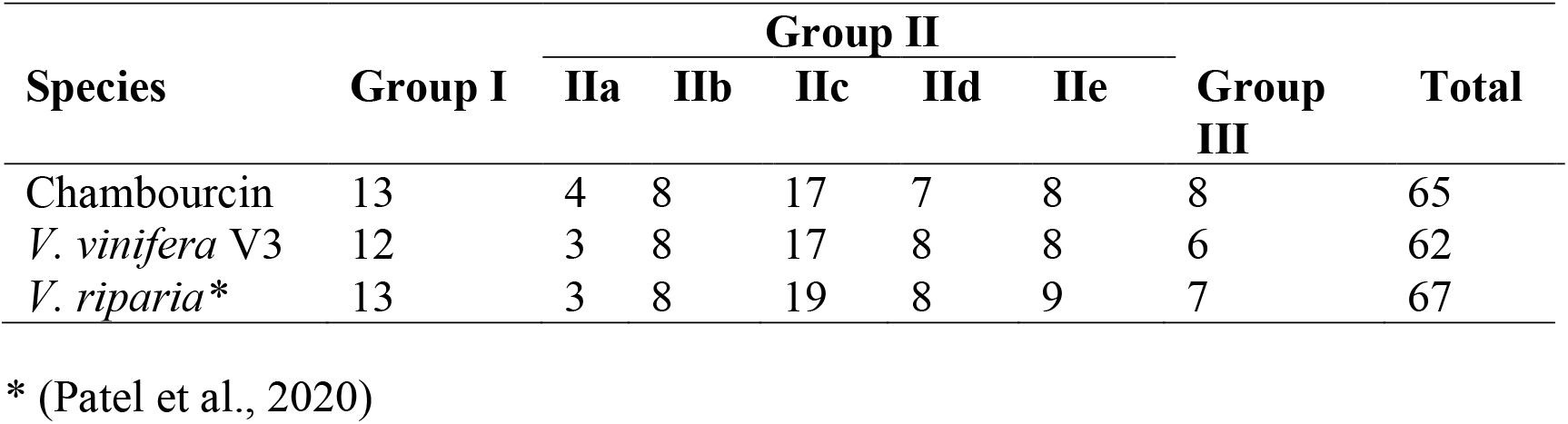
WRKY transcription factor classification comparison of the ‘Chambourcin’ with other grape species.

## Conclusion

In this study, we presented the genome assembly of a complex interspecific hybrid grape cultivar, ‘Chambourcin’, using PacBio HiFi long read sequencing, Bionano third-generation sequencing data and Illumina short read data. The comparative genomic analyses of ‘Chambourcin’ with the reference genome of *V. vinifera* ‘PN40024’ 12X.v2, Shine Muscat and *V. riparia* ‘Gloire’ indicated that the ‘Chambourcin’ genome aligns well to other grape genomes without any large structural variation. Ortholog analyses of ‘Chambourcin’ primary gene models, *V. vinifera* ‘PN40024’ 12X.v2, VCost.v3, Shine Muscat and *V. riparia ‘*Gloire’ revealed that the ‘Chambourcin’ genome assembly and gene annotations are a high-quality grapevine resource for the research community.

Interspecific hybrids derived from two or more *Vitis* species are common in nature (Morales-Crus et al. 2021) and are the cornerstone of grapevine rootstocks grown worldwide, cultivars that predominate in the eastern and midwestern North America, and new disease resistant genotypes currently in development (Migicovksy et al. 2016). The sequence data, scaffold assemblies, and gene annotations of the ‘Chambourcin’ genome assembly described here provide a valuable resource for genome comparisons, functional genomic analysis, and genome-assisted breeding research.

## Supporting information

Table 1 GigaDB

Table 2 GigaDB

Table 3 GigaDB

Table 4 GigaDB

Table 5 GigaDB

## Data Availability

The PacBio HiFi reads and Illumina whole genome reads are deposited in the NCBI BioProject accession PRJNA754438. SRA accession of PacBio HiFi reads is SRR15530464 and SRA accession of Illumina whole genome reads are SRR24093946, SRR24093988, SRR24095403 and SRR24097763. The Bionano maps, genome assembly, gene annotation, proteins and other data are available at figshare: https://doi.org/10.6084/m9.figshare.15505788.v1

## Additional files for GigaDB

**Table 1:** Alignment of ‘Chambourcin’ primary genome assembly with the reference genome *V. vinifera* ‘PN40024’ 12X.v2.

**Table 2.** Mapping of rhAmpSeq with ‘Chambourcin’ genome assembly.

**Table 3.** The Simple Sequence Repeats (SSRs) of the ‘Chambourcin*’* primary genome assembly.

**Table 4.** Functional annotation of the ‘Chambourcin*’* primary proteins.

**Table 5.** The plant transcription factors of the ‘Chambourcin*’* primary and *V. vinifera* ‘PN40024’ 12X.v2, VCost.v3 proteins.

## Author Contributions

S.P., Z.H., A.F. and A.M. conceived of and designed this study, A.F. provided computational resources and guidance for this study and J.P.L. provided ‘Chambourcin’ samples for Illumina whole genome sequencing and rhAmpseq marker haplotypes sequences. S.P. processed DNA sequences for ‘Chambourcin’, assembled the genome and conducted synteny analysis. S.P. processed RNA-Seq data for gene prediction, conducted gene prediction and gene annotation. S.P. conducted comparative genomics analysis, and uploaded sequences to NCBI and figshare; S.P. wrote the first draft of the manuscript. S.P., A.M., Z.H., A.F., and J.P.L. reviewed and finalized the manuscript.

## Acknowledgments

We acknowledge Laszlo Kovacs for collecting ‘Chambourcin’ samples for sequencing and Alex Harkess for assistance developing protocols for DNA extractions from ‘Chambourcin’. Roberto Villegas-Diaz, Chad Julius, Luke Grassman, and Rachael Auch assisted with installing and debugging tools in the South Dakota University Research Cyberinfrastructure High Performance Computing Cluster Roaring Thunder.

